# Eliminating *Aedes aegypti* from its southern margin in Australia: insights from genomic data and simulation modeling

**DOI:** 10.1101/2021.08.21.457232

**Authors:** Gordana Rašić, Igor Filipović, Sean L Wu, Tomás M León, Jared B Bennett, Héctor M Sánchez C, John M Marshall, Brendan J Trewin

## Abstract

A rare example of a successful long-term elimination of the mosquito *Aedes aegypti* is in Brisbane, Queensland, where the legislatively-enforced removal of rainwater tanks drove its disappearance by the mid-1950s. However, a decade-long drought led to the mass installation of rainwater tanks throughout the region, re-introducing critical breeding sites for the mosquito’s persistence in this subtropical region. With *Ae. aegypti* re-invading towns just 150 km north of Brisbane, we examined the potential for their sustained elimination. Through genomic analyses, we estimated historical expansion and current isolation between neighboring populations as close as 15 kilometers. The estimated recent migration rate, entomological and meteorological data were used to calibrate the simulations of elimination campaigns in the two southernmost populations. Our simulations indicate that *Ae. aegypti* could be eliminated with moderate release numbers of incompatible *Wolbachia*-infected (IIT) males (sorted with an error rate ≤10^-6^) if non-compliant rainwater tanks are removed first. With this combined campaign, highly effective suppression (>99%) was predicted in both towns, and complete elimination was predicted in 35% of simulations in one town. Without tank removal, however, IIT led to a moderate suppression (61-93%) even with a 40:1 ratio of released IIT males to local males. Moreover, with a ratio of >20:1, *Wolbachia* establishment was predicted when the sorting error was >10^-7^. Our conservative estimates of intervention outcomes inform the planning of *Ae. aegypti* elimination in the region, and offer insight into the effective combinations of conventional and novel control tools, particularly for vulnerable mosquito populations at range margins.

**Significance:** After decades of range stagnation in Australia, the *Aedes aegypti* mosquito is expanding southward, approaching the most-densely-populated areas of Queensland. Using population genomics and simulation modeling of elimination campaigns, we show that Australia’s southernmost populations of this disease vector are genetically isolated and could be eliminated with moderate releases of incompatible *Wolbachia-infected* males if major larval breeding sites (non-compliant rainwater tanks) are removed first. The risk of *Wolbachia* establishment for this approach is low, and so is the risk of quick mosquito re-invasion. Our conservative estimates of intervention outcomes inform the planning of *Ae. aegypti* elimination in the region, and offer new insight into the benefits of combining conventional and novel control tools, particularly for mosquito populations at range margins.

## Introduction

The globally-invasive mosquito *Aedes aegypti* is commonly known as the “yellow fever mosquito”, although in today’s world it is more important as the major vector of dengue, chikungunya and Zika viruses. Population-genetic data, corroborated by historical records of yellow fever and dengue fever epidemics, indicate that *Ae. aegypti* left Africa some 500 years ago, first invading the Americas, and rapidly expanding across the Asia-Pacific in the second half of the 19th century (1). In Australia, the first reliable record of *Ae. aegypti*’s presence is a museum specimen collected in Brisbane in 1887 (2), but the first local dengue outbreaks occurred earlier and further north, in Townsville in 1879 and Charters Towers and Rockhampton in 1885 (3). Although known as a vector of yellow fever at the time, *Ae. aegypti* was first implicated in the transmission of dengue fever by Thomas Bancroft, while investigating the Brisbane 1905 epidemic (4).

Until the late 1950s, this container-breeding mosquito was widely distributed throughout Australia’s north and west coast, and eastern seaboard for 3,000 km north to south. It is hypothesised that intensive public health interventions after the second World War led to a retraction of its populations into (sub)tropical central and north Queensland (Figure 1) (5). These interventions implemented an approach that relied on intensive surveillance of residential areas to identify key larval mosquito production sources, and enforced compliance by insisting on removal of all water-holding rubbish and noncompliant rainwater tanks (6). The removal or sealing of rainwater tanks is thought to have been a major factor in the elimination of *Ae. aegypti* from the southeast distribution margin, including the city of Brisbane, where they provided key habitat for continual larval development during the cold and dry winter months (6, 7).

**Figure 1.**
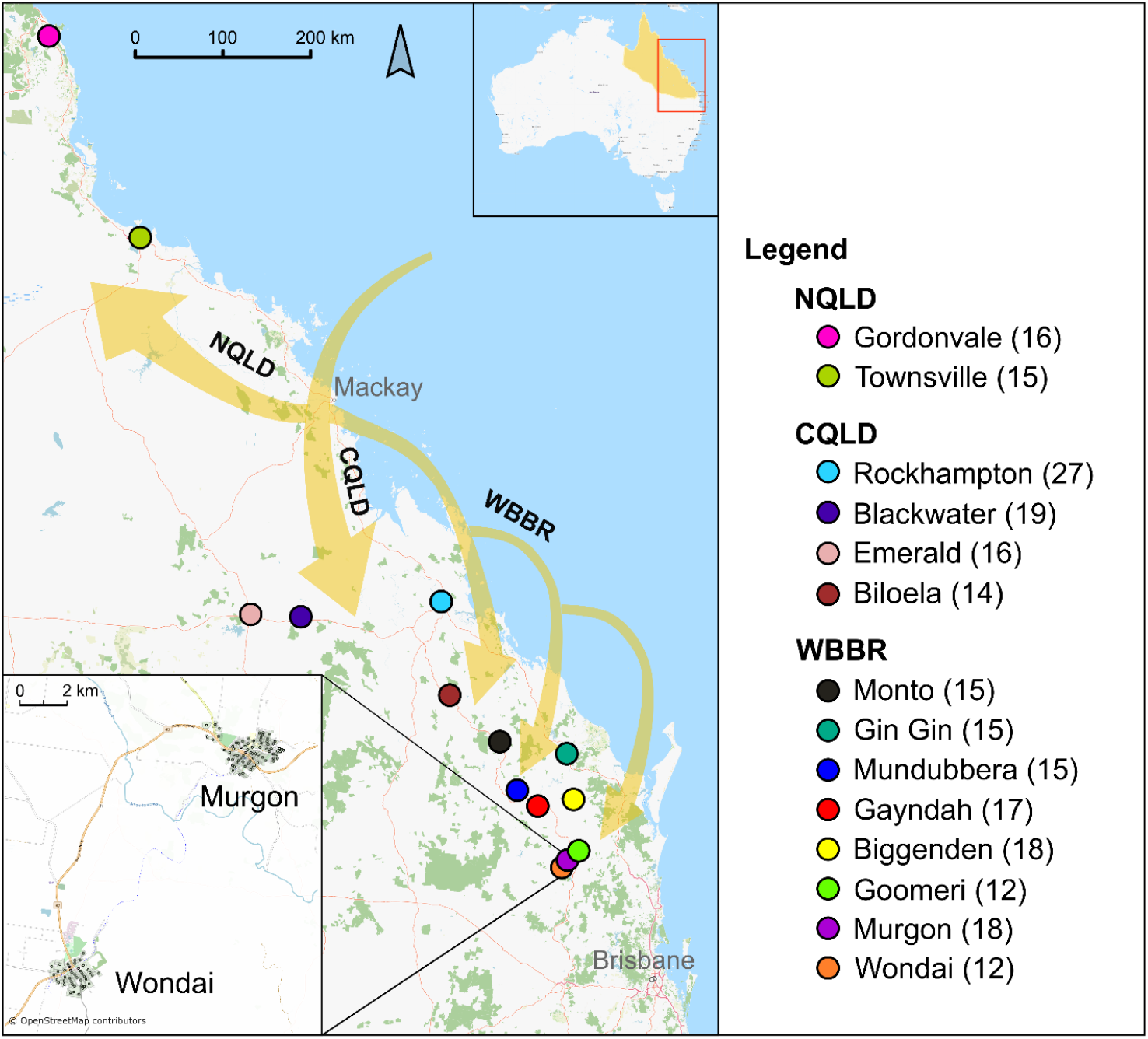
Sampling locations and the inferred population expansion of *Aedes aegypti* in Queensland. Color coding for towns is consistent throughout the manuscript figures. The number of genotyped mosquitoes in each town is shown in brackets. A map in the upper-right corner shows the current distribution of *Ae. aegypti* in Australia (yellow) and the sampled region (red rectangle). A map in the lower-left corner shows Murgon and Wondai with network nodes (centroids of residential blocks) used in the MGDrivE2 simulations. Yellow arrows show the population splitting and historical expansion throughout Queensland (inferred as a maximum likelihood tree from the sampled populations in Treemix, see Figure 2B).

However, the presence of *Ae. aegypti* has been recently confirmed in the southern Wide Bay Burnett Region (WBBR), just 150 km north of Brisbane (Figure 1). Entomological surveys throughout WBBR since 2011 indicate that *Ae. aegypti* populations are larger and more wide-spread along major transport highways than previously thought (Supp. Table S1). It is suspected that *Ae. aegypti’s* re-invasion of WBBR has been facilitated by region-wide installation of rainwater tanks and other changes in water storage practices in response to the region’s driest period on record, the “Millennium Drought”, between 2001 and 2009 (8).

While protected by *Wolbachia* in tropical north Queensland (9), communities in WBBR and central Queensland remain vulnerable to the spread of diseases vectored by *Ae. aegypti* (e.g.(10)). The 2019 dengue outbreak in Rockhampton, the first in over 60 years (11), typifies this vulnerability. The elimination of *Ae. aegypti* populations from the southern margin would not only benefit the towns involved but would also decrease the risk of southward re-incursion into the highly populated areas of southeast Queensland, including metropolitan Brisbane, where there are now over 300,000 rainwater tanks (7).

In theory, the removal of larval habitat would pressure vulnerable mosquito populations through extended dry and cold periods. O’Gower was the first to suggest that *Ae. aegypti* might be eliminated in areas of low rainfall by ‘a continuous mosquito control program, and complete replacement of rainwater tanks by a reticulated water system’ (12). More recent mosquito control activities undertaken in WBBR’s town of Gin Gin (between 2014 and 2019) achieved suppression of *Ae. aegypti* numbers below detection threshold through the sealing of rainwater tanks, a targeted larvicide application to various larval breeding sites, and deployment of lethal adult traps (13). This elimination approach requires extensive community engagement and access to private property for inspection, tank sealing and chemical treatment of mosquito breeding sites by Government-licensed pest management technicians.

In parts of Innisfail in north Queensland, a strong suppression (but not elimination) of *Ae. aegypti* was achieved through releases of incompatible male mosquitoes over 20 weeks during the 2017-2018 field trial (14). This biocontrol strategy, known as the Incompatible Insect Technique (IIT), is analogous to the traditional Sterile Insect Technique (SIT), but instead of releasing male insects sterilized by radiation, the released male mosquitoes carry the bacterium *Wolbachia* that causes death of embryos in matings with wild, or incompatible, females (15). When released in numbers several times greater than the number of local males over the course of several months, incompatible males should effectively render all offspring of local females unviable and eventually crash the population (15). Because IIT males can be released from moving vehicles along public roads (e.g.(16)), this strategy does not require access to private property unless required for entomological monitoring.

IIT could be a viable strategy for eliminating *Ae. aegypti* in WBBR, where most mosquito populations occur in small towns (with <1500 households) separated by an environment inhospitable to *Ae. aegypti*. Production of IIT males and their release across an entire small town, or even two neighbouring towns, should be operationally achievable in WBBR. If effective migration among towns is limited, elimination campaigns could be done successively across the region without much risk of quick mosquito re-invasion from untreated towns.

In this study, we use genomics-informed simulation modeling to assess if IIT and/or the removal of non-compliant rainwater tanks could lead to the local elimination of Australia’s southernmost *Ae. aegypti* populations. We employ a recently developed simulation modeling framework MGDrivE 2 (17), which enables simulations of seasonally-variable mosquito population dynamics and is capable of modeling different control strategies in spatially-structured mosquito populations (17). To explore how vulnerable each WBBR town would be to mosquito re-invasion (post elimination), we ascertained historical and contemporary population connectivity through the analyses of population genomic data from *Ae. aegypti* collected across eight towns in WBBR and six towns located further north (Figure 1). The estimated recent migration rate, along with our entomological surveillance data and meteorological data, were used to calibrate the simulations of elimination campaigns in Murgon and Wondai, the two southernmost *Ae. aegypti* populations (Figure 1).

## Results

### Genomic analyses

The level of historical and contemporary connectivity among *Ae. aegypti* populations in WBBR was estimated using individual-based and population-based genomic analyses. We analysed 229 *Ae. aegypti* individuals collected from 14 localities: eight towns in WBBR (Monto, Gin Gin, Mundubbera, Biggendon, Gayndah, Goomeri, Murgon, Wondai), four towns in central Queensland, CQLD (Rockhampton, Blackwater, Emerald, Biloela), and two in north Queensland, NQLD (Gordonvale, Townsville) (Figure 1). Each mosquito was genotyped across genome-wide SNP positions using the SAM-tools genotype-calling algorithm in ANGSD (18) on the double-digest RAD sequencing data generated with the previously validated protocol (19, 20). The dataset contained 15941 autosomal SNP loci with missingness up to 25% and a depth of ≥5x. Using the same filtering criteria in Stacks v.1.46 (21), we obtained 7716 RAD haplotype loci (Supplemental data).

### Genetic structure and demographic history of *Ae. aegypti* in Australia match geography

Contemporary genetic structure was ascertained using the individual-based analysis implemented in RADpainter and fineRADStructure (22) (Figure 2A). Briefly, the method utilises the information from all SNPs within each RAD haplotype locus, finding one or more closest relatives (nearest neighbours) across RAD loci for each individual and summarizing information into a ‘coancestry matrix’. The clustering algorithm uses a MCMC scheme to find the population configuration (genetic clustering) with the highest probability, and a simple tree-building algorithm produces the posterior probability of assignment to a given population and a hierarchical group (22).

**Figure 2.**
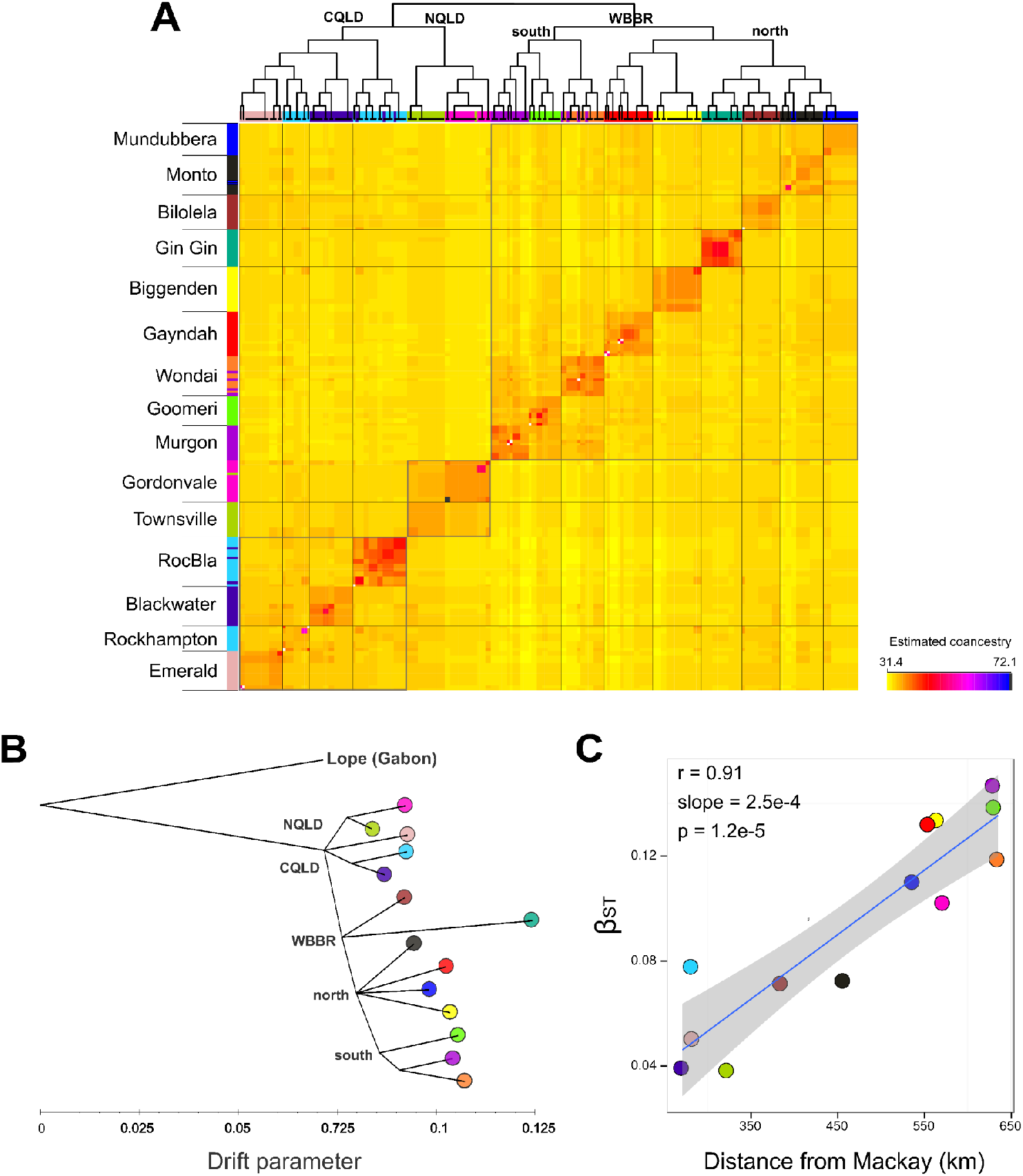
Contemporary genetic structure and historical relationship among *Aedes aegypti* populations in Queensland. (A) Clustered fineRADstructure matrix of coancestry averages per population (B) A rooted maximum likelihood tree of populations (consensus tree from 1000 Treemix analyses, all branches have ≥90% support). (C) Correlation between the population-specific differentiation and geographic distance from the hypothesized ancestral location (port of Mackay).

The results show that *Ae. aegypti* from the same town share more coancestry with each other than with mosquitoes from different towns. In other words, every town contains a genetically distinct population and these genetic groups (clusters) are visible as orange-red squares across the diagonal of the coancestry matrix (Figure 2A). There are some exceptions: a few individuals sampled in Murgon are grouped with individuals from neighboring Wondai, and there was a cluster of individuals from Rockhampton and Blackwater, representing a family group. These cases indicate recent migrants.

The hierarchical structure among populations is clearly inferred (with all posterior probabilities >0.95, Suppl Figure S1), visualized as a tree at the top of the coancestry matrix (Figure 2A). Geographic region defines the highest hierarchical grouping, with a lineage of populations in central Queensland (CQLD), another lineage in north Queensland (NQLD), and the WBBR lineage that splits into the north-central group and the southern group of populations (Goomeri, Murgon, and Wondai). While falling under the jurisdiction of CQLD, Biloela is geographically close to the WBBR border and its *Ae. aegypti* population is more related to the WBBR lineage than to the CQLD lineage (Figure 2A).

To distinguish older and more recent splitting between identified populations, we used Treemix (23) that infers the topology of historical relationships among populations, under the assumption that their history is approximately tree-like. This analysis uses population SNP frequencies and an approximate model for genetic drift to resolve the pattern of population splits and mixtures, where contemporary populations are related to a common ancestor *via* an ancestral population graph (23). We rooted the population tree with a sample from *Ae. aegypti’s* ancestral range, La Lope in Gabon (24). The resulting consensus population tree (Figure 2B) matches the hierarchical clustering inferred with fineRADstructure: the earliest population splitting corresponds to the three geographic regions - NQLD, CQLD and WBBR. The population in Biloela, a town situated near the CQLD-WBBR border, is more related to the WBBR lineage than to the CQLD lineage. In WBBR, the pattern of ‘star-like’ branching in north-central populations (Monto, Mundubbera, Biggenden, Gayndah) indicates mass expansion from a common source. A stepping-stone southward expansion into Goomeri, Murgon and Wondai formed another lineage (Figure 2B). The population in Gin Gin has undergone a substantial genetic drift, representing a genetic outlier in WBBR. This is not surprising, as we sampled *Ae. aegypti* in this town only one year prior to the local elimination (i.e. at the very tail of the public health campaign).

### Serial north-to-south expansion into WBBR is not from Rockhampton

The inferred historical relationships among *Ae. aegypti* populations support the hypothesized north-to-south expansion into the WBBR. Rockhampton was first hypothesized as a source for the linear expansion into WBBR by Kay *et al*. (5) based on intermittent *Ae. aegypti* surveys between 1956 and 1983. However, our genomic analyses indicate that the WBBR lineage and CQLD lineage (that includes Rockhampton) originate from the same ancestral region.

One method to infer the direction of expansion from one ancestral source involves finding a correlation between the population-specific genetic parameter *β*st (25) and geographic distance from the suspected ancestral location (e.g. (26)). The expectation is a positive correlation between these two parameters – i.e. *β*st should be higher for populations that are farther from the ancestral source. Also, βst is expected to be negative for populations in or near the ancestral location (25). None of the analysed populations had a negative *β*st, indicating that we did not sample population(s) closest to the ancestral source, but the smallest value (*β*st = 0.04) was detected for Townsville (NQLD) and Blackwater (CQLD) (Figure 2C). Assuming the population origin on the coast between NQLD and CQLD, we considered Mackay, the first port of entry for European settlers on the Australian tropical coastlands (27), and found a highly significant positive correlation between *β*st and distance from Mackay (*r* = 0.91, *p* = 1.2e-5, Figure 2C). The correlation was smaller when distance from Townsville was used (*r* = 0.77, *p* = 0.005) and non-significant if distance from Blackwater was used (*r* = 0.18, *p* = 0.563). The highest *β*st were recorded for populations at the most southern margin, 0.14 and 0.15, for Goomeri and Murgon, respectively. (Note that we excluded Gin Gin from this analysis as a demographic outlier with *β*st of 0.24).

This is the first insight into the demographic history of *Ae. aegypti* in Australia, with multiple analyses indicating initial establishment between north and central Queensland and expansion in multiple directions, including a series of southward introductions into WBBR.

### Most WBBR populations have low effective population size and high isolation

Excluding Gin Gin as a demographic outlier, genetic diversity of *Ae. aegypti* populations in WBBR decreased along the north-south direction, with significant correlation between latitude and the expected heterozygosity (*r* = 0.885, *p* = 0.008). The effective population size (*N_e_*) exhibited the same pattern, ranging between 21 and 120, but the most southern population in Wondai represented an outlier with *N_e_* that is 3-17 times greater than in other WBBR populations (Figure 3A). For *Ae. aegypti* collected in Gin Gin in 2016, *N_e_* was only 4 (95% CI: 3-4), clearly reflecting the effects of the public health campaign (2012-2016) that has led to undetectable mosquito populations from 2018 onwards (13). The entomological survey in 2018 recorded 29-100% of inspected premises positive for adult *Ae. aegypti* (Figure 3A), and 20-100% positive ovitraps in WBBR towns (Table S1). Interestingly, we detected a strong correlation between the percent of positive households for adults (entomological index reflecting the population abundance) and *N_e_* (genetic parameter reflecting, among other processes, the variability in abundance (28, 29)), with *r* = 0.945, *p* = 0.001 (excluding Wondai as an outlier). This suggests use of effective population size as a proxy for how many households have adult *Ae. aegypti*, avoiding extensive and costly house-to-house surveys. Outliers like Wondai, however, indicate that more data are needed to robustly ascertain the predictive value of this relationship.

**Figure 3.**
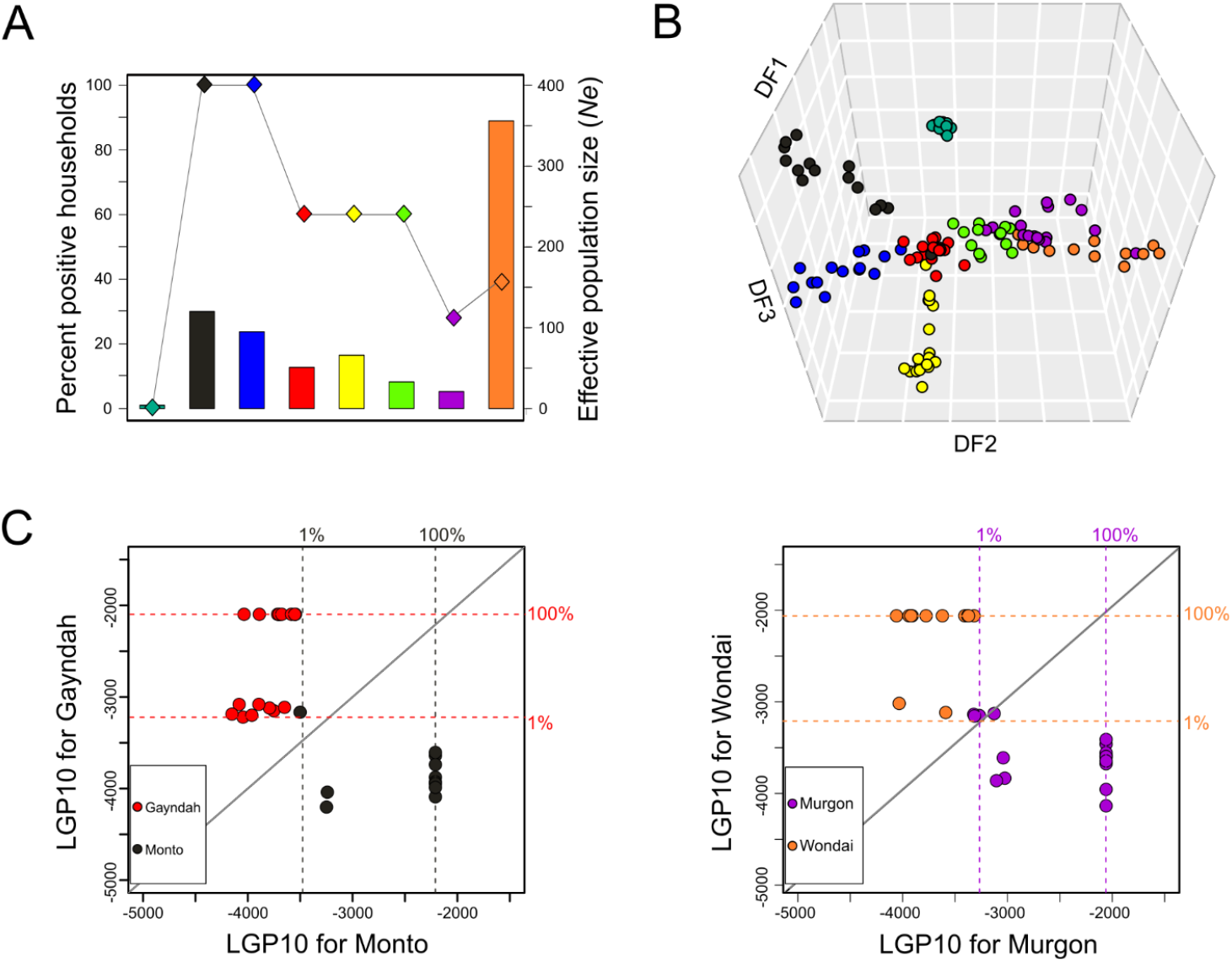
Effective population size and genetic isolation of *Aedes aegypti* populations in WBBR. (A) Effective population size (*N_e_*) and the percent of positive households for adult *Ae. aegypti* in eight towns in WBBR. (B) The first three axes from the DAPC with eight genetic groups in WBBR. (C) Genetic assignment analysis in GenePlot for detection of recent migrants between Monto and Gayndah (left panel), and Murgon and Wondai (right panel). Each individual is presented as a point that is color-coded according to the sampling location (e.g. black if sampled in Monto, red if sampled in Gayndah). 1% and 100% quantiles of the distribution of log genotype probabilities (LGPs) for each population are represented with dotted lines.

Genetic structuring in WBBR closely matches geography - the Discriminant Analysis of Principal components (DAPC) (30) is able to group together individuals sampled in the same town (represented as points, color-coded based on the sampling location, Figure 3B). This multivariate analysis produced a clear separation of town samples (genetic clusters) across the first three axes: the first axis (DF1) separates Gin Gin from other populations, the second (DF2) separates the north-central from the southern populations, and the third (DF3) further separates populations within the north-central WBBR.

We used the genetic assignment visualization method in GenePlot (31) to ascertain the individual’s absolute genetic fit to the inferred populations, and therefore to detect recent migrants as individuals with a good fit to populations other than where they were sampled. We found that one mosquito sampled in Monto showed a good fit to the Gayndah population (black point within red dotted lines, Figure 3C), indicating a putative first-generation migrant - i.e. an individual that was collected in Monto but has both parents from Gayndah. An individual that shows an equally good fit to both populations (its LGP falls on the thick diagonal line in the plot), represents a putative second-generation migrant, with one parent from each population. In Murgon, we detected one putative second-generation migrant and two putative first-generation migrants (Figure 3C), indicating a substantial recent gene flow from Wondai to the neighbouring Murgon. Given that female *Ae. aegypti’s* active flight range and intergenerational dispersal rarely exceed 200 meters (32, 33), the distance between towns in WBBR (at least 15 km) is certainly traversed passively through human transport such as commuting, cargo shipping, etc. Recent migrants were not detected in any other GenePlot analysis of population pairs, indicating high genetic isolation of most WBBR populations.

With strong genomic evidence for southward expansion and contemporary migration between the two southernmost WBBR populations, we simulated the elimination programs that would first simultaneously target *Ae. aegypti* in Murgon and Wondai.

### Simulation of elimination campaigns

Using MGDrivE 2 (17), we simulated landscapes in Murgon and Wondai as metapopulation networks, where each node in the network represents a housing block, and blocks have a median of 17 houses. Each household was modeled to have ~6 adult female *Ae. aegypti*, based on entomological surveillance data (see Materials and Methods), and seasonal variation in *Ae. aegypti* population density was driven by local temperature and rainfall data, which determine adult mortality rates, juvenile developmental rates and larval carrying capacity. Daily dispersal of adult mosquitoes between nodes was calibrated to previously-generated mark-release-recapture data (34), and batch migration between towns was inferred from our genetic assignment results (see Materials and Methods). For each of the modeled elimination strategies, we ran 100 stochastic simulations for 5 years each in order to account for variation in the stochastic model and chance events such as population elimination. The modeled elimination strategies include:

1. Removal of non-compliant rainwater tanks at the beginning of year 3 (Figure 4A),
2. Releases of IIT males at the beginning of year 3 - twice a week for 12 weeks, with each release containing 14-40 times the number of males present in year 1 ( the “overflooding ratio” of 14:1, 20:1, 30:1, 40:1) (Figures 4B and 4D), and
3. A combined strategy, with the removal of non-compliant rainwater tanks at the beginning of year 3, and IIT releases at the beginning of year 4 - twice a week for 12 weeks with an overflowing ratio of 14:1 (as compared to the number of males present during the peak season in year 1) (Figure 4C).

**Figure 4.**
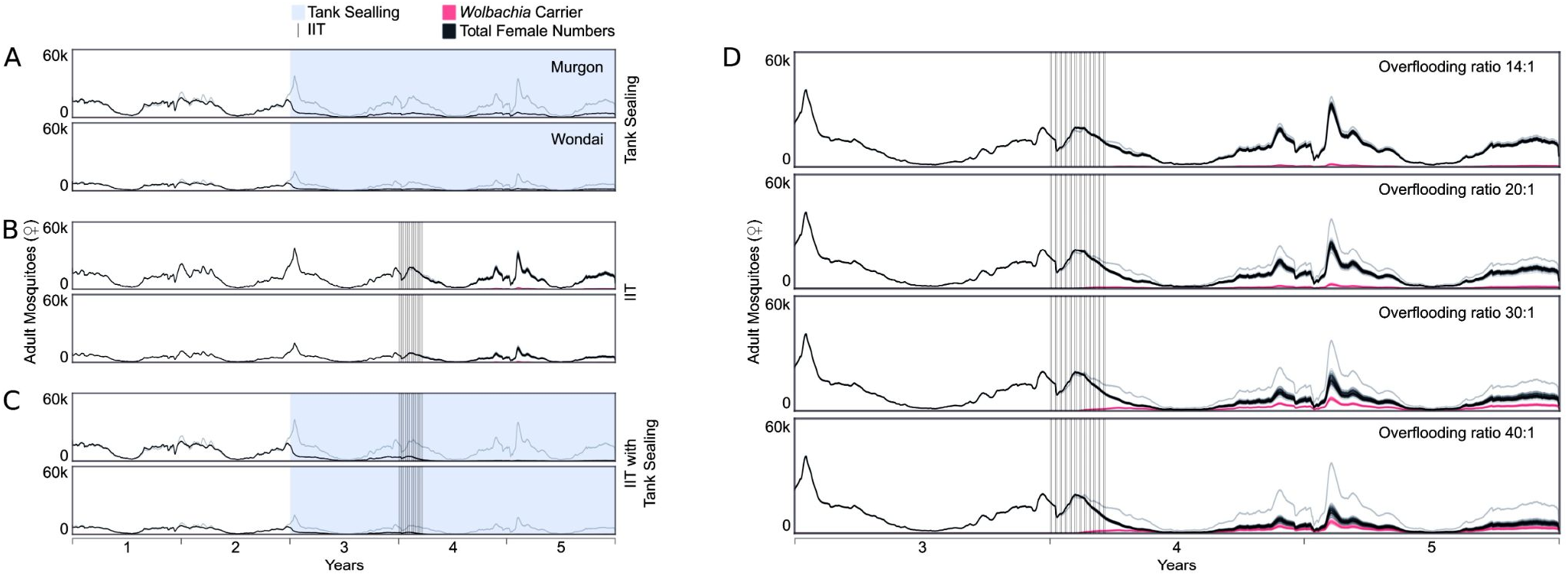
MGDrivE 2 simulations of tank sealing (source reduction), IIT (releases of *Wolbachia-carrying* males) and the combination of these two interventions in settings resembling Murgon and Wondai. Each setting is modeled as a metapopulation of residential blocks, and results represent the mean of 100 stochastic simulation runs. Seasonal variation in adult *Ae. aegypti* population density is driven by local temperature and rainfall data, which are used to calculate time-varying adult mosquito mortality rates, juvenile development rates and larval carrying capacity, as described in the Materials and Methods. Mean *Ae. aegypti* life history parameters are listed in Table S2. The resulting total adult female population size in Murgon and Wondai is denoted by black lines, and faint grey lines in the absence of interventions. (A) Tank sealing is modeled in Murgon and Wondai, starting 2 years into the simulation and denoted by the shaded light blue portion of the plot. One sixth of the maximum larval carrying capacity in each town is assumed to be provided by noncompliant rainwater tanks that may be sealed. (B) IIT releases are modeled in Murgon and Wondai, starting 3 years into the simulation and occurring twice a week for 12 weeks, with each release (denoted by vertical black lines) containing 14 times the number of males as present in year 1 (the “overflooding ratio”). Due to imperfect sex-sorting, *Wolbachia-carrying* females are included in the IIT releases at a frequency of 10^-6^. This leads to *Wolbachia-carrying* females emerging and persisting in each setting, denoted by magenta lines. (C) The tank sealing intervention is combined with IIT releases at an overflooding ratio of 14:1. (D) IIT releases without tank sealing are modeled in Murgon for overflooding ratios of 14:1, 20:1, 30:1 and 40:1. Results suggest that a combined intervention of tank sealing with IIT releases could lead to local *Ae. aegypti* elimination for a moderate overflooding ratio of 14:1 (panel C). Neither intervention leads to elimination on its own (panels A and B), and larger IIT releases lead to establishment of *Wolbachia-carrying* females (panel D), hindering the impact of subsequent releases.

Simulations were continued for three years after the start of the tank sealing intervention in scenario 1 and 3, and for two years after the start IIT releases in scenarios 2 and 3. We assessed the effectiveness of each scenario by calculating (i) the proportion of simulations with zero adult females across blocks over the entirety of year 5, and (ii) the reduction in the total number of females per day, relative to simulations without any intervention, averaged across simulation runs.

For each of the three scenarios (Figure 4A-C), the total adult female population size over 100 simulation runs is shown as solid black lines, while faint grey lines represent female population size without any intervention. Seasonal fluctuation in both towns is very pronounced, with adult numbers peaking during summer (January-March) and reaching only 8.6% and 9.2.% of the peak size (in Wondai and Murgon, respectively) during the cold winter months (June-August). With the removal of non-compliant rainwater tanks (Figure 4A), reduction in adult female numbers occurs quickly and persists through the following years, suppressing the population to 24% that of the non-treated population in each town. With 12 consecutive weeks of moderate IIT releases (overflooding ratio of 14:1, Figure 4B), a small reduction in mosquito numbers was predicted the following year: 11.6% suppression in Murgon and 15.6% in Wondai (Figure 4B). A significant population reduction (but not elimination) is only predicted with the highest overflooding ratio (40:1) and is comparable in impact to the tank-sealing campaign (~78% suppression, Figure 4D). Moreover, when the rate of contamination with *Wolbachia-carrying* females (due to imperfect sex sorting) is greater than 10^-7^, IIT with an overflooding ratio of 20:1 or more consistently leads to *Wolbachia* establishment (as shown by the solid magenta lines in Figures 4B and 4D).

Our simulations show that *Ae. aegypt*i could be eliminated by implementing a combined strategy, whereby a moderate IIT regime (overflooding ratio of 14:1) is conducted a year after the tank-sealing campaign (Figure 4C). In Wondai, complete elimination (zero females across all residential blocks over the entirety of year 5) was predicted in 35% of simulations, and in the remaining 65% of simulations local elimination was predicted in 95-98% of residential blocks. The total number of adult females per day in Wondai was on average 186 times higher in simulations without any intervention, which is equivalent to the average population suppression of 99.5%. Complete elimination was predicted in only 4% of simulations in Murgon, but local elimination was consistently predicted in >88% of residential blocks. The daily number of adult females in Murgon was suppressed by 99.2% (Figure 4C). For this combined strategy, *Wolbachia* establishment was not predicted even with the highest sex sorting error we simulated (10^-6^, Figure 4C).

## Discussion

Mosquito elimination programs require a comprehensive effort supported by sustained funding, experienced staff and community participation, as well as extended monitoring activities to confirm mosquito absence (35). Very few successful attempts to eliminate established populations of *Ae. aegypti* through conventional programs have been reported to date. Such examples include the campaigns across South America between 1940s and 1970s, and areas of Cuba in the 1980s that employed intensive insecticidal treatment and source reduction, but these programs were unsustainable (36). The only long-term successful elimination of this disease vector appears to be in Brisbane, Queensland, where the legislatively-enforced removal of rainwater tanks drove *Ae. aegypti’s* disappearance by the mid 1950s (6). However, a decade-long drought during the 2000s led to the installation of hundreds of thousands of rainwater tanks throughout south-east Queensland, including Brisbane, providing potential critical larval habitat for *Ae. aegypti* persistence in this subtropical region (6). With the discovery of *Ae. aegypti* populations in towns just 150 km north of Brisbane (Figure 1, south Wide Bay Burnett Region), we examined the potential for their elimination with accessible vector control tools.

Our entomological survey in the WBBR indicates that *Ae. aegypti* populations are larger than anticipated, but tend to decrease in southern towns (Figure 3A). Our genomic analyses support the hypothesis of *Ae. aegypti* serial north-to-south expansion into the WBBR (Figure 2B-C), and indicate that populations are analogous to isolated island populations (Figure 3B-C), which is favorable for achieving long-term elimination. A recent public health control program in the WBBR town of Gin Gin provides empirical evidence for our genomic-based inference. Namely, *Ae. aegypti* numbers in this town have remained below detectable levels three years after the completion of the public health mosquito control program (13). Given the high level of genetic structuring in WBBR where each population has a specific genetic profile, ongoing monitoring should be able to distinguish between incomplete elimination and re-invasion of *Ae. aegypti* individuals from outside populations. Genomic data have been used to track *Aedes* invasions (e.g. (37–39)), including the recently established populations of *Ae. aegypti* in California (40, 41). However, our study is the first to demonstrate remarkable power in ascertaining specific genetic profiles of invasive mosquito populations from neighboring towns as close as 15 kilometers (Figures 3B-C).

In addition to creating baseline population data for monitoring purposes, high-resolution genomics enabled us to quantify recent mosquito migration between Murgon and Wondai (genetic assignment test, Figure 3C). We used this genomically-inferred migration rate to parametrize batch migration between towns in our spatially-explicit simulations of elimination campaigns (MGDrivE2, Figure 4). To parameterize fine-scale dispersal rate (i.e. movement of *Ae. aegypti* between residential blocks within a town), we used the results of a mark-release-recapture (MRR) experiment in one of the WBBR towns (34). However, Filipovic *et al*. (32) recently demonstrated that the analyses of fine-scale spatial genomic data, such as separation distance between close kin, can be a powerful alternative to MRR (32). Increasingly, high-resolution genomics is becoming a powerful tool to support mosquito control modeling, monitoring and decision making.

Simulation modeling of current and historical *Ae. aegypti* persistence across Australia predicts high natural vulnerability of subtropical populations (such as those in WBBR) due to suboptimal climate (42, 43). However, this vulnerability depends strongly on the availability of large larval breeding sites like rainwater tanks (7, 42). The removal/sealing of non-compliant rainwater tanks should therefore significantly increase the probability of *Ae. aegypti* elimination in WBBR. A public health campaign in the WBBR town of Goomeri in 2018, which primarily focused on the inspection and sealing of rainwater tanks, led to the suppression of *Ae. aegypti* in the following year, with 75-80% fewer females caught during the peak productivity season (February-March 2019, File S1). While sealing of non-compliant tanks seems insufficient to achieve elimination of *Ae. aegypti* on its own, our simulations show that its implementation ahead of the IIT campaign is critical (Figure 4). Namely, our simulated releases of *Wolbachia-infected* (IIT) males led to a moderate population suppression (~78% reduction in female *Ae. aegypti* numbers) even with a 40:1 overflooding ratio of IIT males to wild-type males (Figure 4B). However, when tank sealing was simulated one year prior to the IIT releases, highly effective suppression (>99% reduction in adult female *Ae. aegypti* numbers) was observed in both towns, and the complete elimination was observed in 35% of simulations in one town (Figure 4C)

A recent IIT field trial in Innisfail, northern Queensland, achieved high suppression (>80%) across three treatment sites, with releases conducted three times a week over 20 weeks and an overflooding ratio of ~10:1 (14). Our simulations of the releases conducted twice a week over 12 weeks predict that increasing the overflowing ratio from 14:1 up to 40:1 achieves suppression levels 61-93%, but larger releases (≥20:1) dramatically increase the chance of *Wolbachia* establishment if the sex sorting error is greater than 10^-7^ (Figure S1). This is consistent with a recent modeling study of the IIT campaign in Innisfail (44) which found that a sex sorting error rate of 10^-7^ or less is needed to achieve a low probability of *Wolbachia* establishment, along with ‘adaptive releases’ three times a week that start with an overflooding ratio of 5:1 or 15:1, and decrease in sze as the population is suppressed over time (44).

Depending on the underlying technology, the reported sex sorting error in IIT programs is between 10^-3^ and 10^-9^ (16, 45), and we opted to simulate the releases with the sorting error rates from 10^-6^ to 10^-9^ (File S1). It is worth noting that our simulations could be overestimating the risk of *Wolbachia* establishment under imperfect sex sorting, as they assumed that *Wolbachia* infection had no fitness cost. While the laboratory experiments revealed that transfections with *Wolbachia* strains like *w*AlbB and wMel had no significant cost for *Ae. aegypti* males (46, 47), *Wolbachia* decreased female fecundity and survival, as well as hatching of eggs that were quiescent in warm environments (48). However, field data from recent IIT trials in Singapore reported *Wolbachia* (*w*AlbB) establishment in some treatment areas with an overflooding ratio of 30:1, even with the deployment of a sex-sorting technology with the highest fidelity (45). One mitigation strategy for this issue is the irradiation of IIT mosquitoes (SIT-IIT) to ensure that accidentally-released *Wolbachia-carrying* females are also sterile (49). However, irradiated IIT males are less competitive than non-irradiated males (45) and thus necessitating higher release ratio and/or frequency. Another approach to minimize the chance of *Wolbachia* establishment could be to use a *Wolbachia* strain with high fitness cost like *w*MelPop (50). Such a strain is difficult to establish even with very high numbers of released *Wolbachia-carrying* females (51), and if established, would likely cause a population crash during dry winter months in WBBR because it causes high egg mortality under such conditions (52, 53). The release ratio and frequency for the *w*MelPop-carrying males, like for the irradiated IIT males, would have to be high. Conversely, our simulations indicate that *Ae. aegypti* populations in WBBR could be eliminated with a moderate release frequency and numbers of IIT males sorted with moderate fidelity (1-10^-6^), if non-compliant rainwater tanks are removed first (Figure 4C).

The importance of simulation modeling to predict the impact of integrated mosquito control programs tailored to specific ecological settings has been widely recognized in malaria elimination campaigns (54). To our knowledge this is the first modeling analysis of an integrated elimination strategy, combining source reduction and IIT against the dengue vector, *Ae. aegypti*. We considered the scenarios where rainwater tank sealing and the IIT campaign both start at the peak season of mosquito productivity. However, in regions like WBBR where mosquito populations experience highly seasonal dynamics, starting both interventions ahead of the peak season is expected to improve the effectiveness of a campaign. Finding solutions that maximize population suppression could be further tackled through the use of optimization models for vector control (54). Our estimates of intervention outcomes are conservative, informing a robust public health plan for the elimination of *Ae. aegypti* in the WBBR.

Here we demonstrate the benefits of combining larval source reduction with a novel technology like IIT to eliminate recently-established *Ae. aegypti* populations at the range margin in Australia. These benefits are also likely to exist for similar populations in other parts of the world. For example, in less than a decade since its first detection in California (55), *Ae. aegypti* has established large populations that have proven difficult to suppress, as evidenced by the recent IIT trials in Fresno County (16). During the same period, the state has been experiencing major droughts (56), which led to changes in water harvesting and storage policy, such as the introduction of the Rainwater Capture Act of 2012 (57), incentivizing the installation of rainwater tanks on residential properties. Incorporating the sealing/ removal of non-compliant water tanks into the mosquito control programs that utilize IIT could prove critical for the elimination of *Ae. aegypti* populations in California as well.

## Materials and Methods

### Mosquito sampling

*Aedes aegypti* were surveyed and sampled across towns in the Wide Bay Burnett Region (WBBR) between February and April 2018, except in Gin Gin where the sample was collected in 2016 (the last year *Ae. aegypti* was detected in this town). Towns in central Queensland (CQLD): were surveyed and sampled from January to April 2019. The samples from north Queensland (NQLD) were described in two previous studies: *Ae. aegypti* from Gordonvale was collected in 2010 (19), and from Townsville in January 2014 (58).Importantly, these NQLD samples were collected prior to the enrollment of the *Wolbachia* replacement program in the region (59, 60). In the WBRR, trapping was done across five premises, with one BGS trap and four ovitraps per premise, inspected weekly. Eggs were dried and stored in cooler bags, before being reared within until the 4th instar stage. In CQLD, adults were collected by the public health officials using Gravid Aedes Traps (GAT), and 4th instar larvae during house-to-house surveys. Adults and larvae were identified to species (61) and *Ae. aegypti* were stored in 90% ethanol at −20°C. To avoid the sampling of highly related individuals, we selected a single adult or larva from each trap at different collection dates where possible.

### Mosquito genotyping

Total genomic DNA was extracted from individual adults and larvae using DNeasy Blood and Tissue DNA extraction kit (Qiagen, Hilden, Germany), quantified with the Qubit High Sensitivity DNA kit (Thermo Fisher Scientific, Waltham, MA, USA) and 100 ng used for downstream processing. Double-digest Restriction site Associated DNA sequencing (ddRAD-seq, (20)) was used to generate genome-wide SNP and haplotype data for 229 *Ae. aegypti* individuals, following the previously described protocol for library preparation and sequence data processing (19). We used the SAMtools genotype calling algorithm in ANGSD (18).

### Inference of population structure and history

To infer contemporary population genetic structure, we used the individual-based analysis implemented in RADpainter and fineRADStructure (22) with RAD haplotype loci generated in Stacks v.1.46. The input file for RADpainter was created using the script ‘Stacks2fineRAD.py’ that processes the output format of the Stacks *populations* program (‘batch_1.haplotypes.tsv’) (62). First, the RADpainter program utilises the information from all SNPs within each RAD haplotype locus to find the most recent coalescence (common ancestry) among the sampled individuals. Second, the fineRADstructure program uses a MCMC scheme to cluster the individuals, retaining the population configuration with the highest probability. A visualization of the clustered coancestry matrix enables easy inference of the number of genetic clusters, quantification of ancestry sources in each group, and inference of relationships between groups through a simple tree-building algorithm (22).

Historical relationship among populations was inferred in Treemix (23) which produces a population tree under a model where contemporary populations are related to a common ancestor *via* a graph of ancestral populations. This tree is optimal for inferring the topology of relationships among populations, rather than the precise timing of demographic events (23). Using the frequencies of genome-wide SNPs from population samples, the pattern of population splits and mixtures is inferred through a Gaussian approximation to genetic drift. To infer the topology, we rooted the tree using a sample from La Lope, Gabon as an outgroup, given that this population resides in *Ae. aegypti’s* ancestral range (24). We filtered 9954 SNPs that were found in the African and all Australian population samples, and used a window size of 50 (app. 2.5 Mb-blocks with 50 SNPs) to account for linkage between nearby SNPs. The analysis was repeated 1000 times to produce a consensus tree using the 90% support threshold.

Genetic clustering within WBBR was done using Discriminant Analysis of Principal Components (DAPC) (30), implemented in the R package adegenet v.1.42 (63). DAPC finds linear combinations of alleles (also known as the discriminant functions, DFs) which best separate genetic clusters. We used the genetic clusters that were inferred in fineRADstructure, as well as *a priori* groups based on their sampling origin (towns). The R package plot3D (64) was used to visualize the separation of clusters along the first three DFs.

To infer the direction of expansion into the WBBR, we first aimed to identify populations at or near the ancestral/source location, using the recently developed population-specific genetic parameter *β*st (25). For a set of populations, those with the smallest *β*st should be the closest to the ancestral population (e.g. (26, 65)). Negative *β*st is expected for large ancestral populations that have had time to accumulate many private alleles at low to intermediate frequencies (25). A linear increase in *β*st with the distance from the ancestral location is expected under the isolation-by-distance. Genome-wide *β*st and standard error were estimated using the R package FinePop2 v.0.2 (65), and the lm() function was used to estimate the slope of the linear relationship between *β*st and the Euclidean distance (in km) from the likely ancestral location.

### Inference of effective population size and recent migration

Contemporary effective population size (*N_e_*) was estimated for each WBBR population, using the linkage disequilibrium-based method by Waples *et al*. (66) as implemented in the program NeEstimator v.2.1 (67). Each population had a different number of monomorphic loci and singleton alleles that were excluded from the total of 15941 input loci, producing the estimates with 11708 to 13274 loci across eight populations.

We used the genetic assignment method in GenePlot (31) to identify recent migration events between WBBR populations. The method calculates the absolute fit of each individual to a given population, and the assignment results are presented graphically as log genotype probability (LGP) plots with graph positions for each individual. Every genotyped individual has some missing loci (commonplace for RADseq data), and GenePlot calculates appropriate plot positions for incomplete genotype profiles (31). Recent migrants are identified as individuals that do not have a good fit for the population where they were sampled from - i.e. have low LGP that falls outside the 1% quantile of the LGP distribution for the population of their sampling.

### Simulations in MGDrivE 2

To model the expected performance of tank-sealing and IIT releases at suppressing and eliminating local *Ae. aegypti* populations in Murgon and Wondai, we used the MGDrivE 2 framework (17). This framework models egg, larval, pupal and adult mosquito life stages with overlapping generations, and larval mortality increasing with larval density. The inheritance pattern of *Wolbachia*, resulting from maternal transmission and cytoplasmic incompatibility, was modeled within the inheritance module of the framework. No fitness costs were assumed for *Wolbachia-carriers*, which was a conservative estimate for modeling the risk of establishment during an IIT program. We distributed *Ae. aegypti* populations according to housing blocks, with a median of 17 houses per block, and block centroid coordinates calculated in ArcGIS. Each household was modeled with ~6 adult female *Ae. aegypti* during the peak season of year 1, based on the entomological surveillance data (File S1).

Seasonal variation in *Ae. aegypti* population density was driven by local temperature and rainfall data, which determine adult mortality rates, juvenile development rates and larval carrying capacity (17). For the 5-year simulations, we used data for temperature and rainfall from the Australian Bureau of Meteorology for Murgon and Wondai for the years 2009 through 2013. The hourly temperature data were obtained from the weather station closest to Wondai (30 km south, Kingaroy Airport weather station; lat: 26.5737, lon: 151.8398). Daily rainfall data were obtained from the PostOffice station records in Murgon and Wondai. Temperature-dependent adult mortality and juvenile developmental rates were derived from the functions for *Ae. aegypti* from Mordecai *et al*. (68), and daily larval carrying capacity was derived using the method from White *et al*. (69) in which carrying capacity is proportional to past rainfall weighted by an exponential distribution with a mean of eight days. We calibrated static expected larval carrying capacities for each town based on non-time-varying simulations with fixed parameters, and then rescaled these values as rainfall-dependent time series using the static values as means. We further assumed that ⅙ of maximum larval carrying capacity in each town was provided by non-compliant rainwater tanks (permanent larval breeding sites), and the remainder by recent rainfall that fills up small breeding containers. This composition was based on the surveillance data from Goomeri to match their reduction in mosquito numbers observed one year after the rainwater tank-sealing campaign in 2018 (File S1).

.Active mosquito movement between nodes was parameterized based on a mark-release-recapture experiment in Gin Gin (34), assuming that each adult mosquito has a 1% lifetime probability of movement to any neighboring nide (housing blocks sharing a border), and with each potential destination node being assigned equal probabilities. The probabilities were converted into daily rates for the simulation model. Batch migration of immature and adult mosquitoes (passive migration) between Wondai and Murgon was parameterized using the genomics-inferred frequency of recent migrants (described below). MGDrivE 2 was implemented as a stochastic simulation model to capture random effects at low population sizes and the potential for population elimination. For each of the modeled elimination strategies, 100 stochastic simulations were run, and adult female mosquito counts (including those carrying *Wolbachia*) were recorded. Average mosquito life-cycle parameters used in these simulations are provided in Table S2.

#### Dispersal and batch migration parametrization

We assumed each mosquito (both sexes) had a 1% lifetime probability to leave their natal node (block) and migrate to a neighboring node (housing blocks sharing a border), and choice of specific destination node were given uniform probabilities. Because adult mosquito mortality is temperature dependent, we used the yearly average to calculate mean mosquito lifespan, from which we obtained the dispersal rate such that the probability of leaving the natal node before death was 1%.

We parameterized batch migration of egg and adult stages from Wondai to Murgon using the genetic assignment data. Genetic assignment revealed 2 individuals in Murgon for whom both parents had originated in Wondai, indicating these individuals were direct migrants, and an additional individual with mixed parentage (Wondai/Murgon) indicating it was the offspring of a direct migrant. Assuming the sample was unbiased, ~10% of the population in Murgon are direct migrants from Wondai. Using the estimate of 14,268 for the total mosquito population size in Murgon, the population who are not direct migrants is therefore 12,841. Of the remaining mosquitoes in the sample, we assume that they were offspring of unique mating pairs. To estimate the number of these parents which were from Wondai, assuming mosquitoes are randomly mating and that they were offspring of previous direct migrants (not direct migrants in the same sample), the number of parents from Wondai from sampled (without replacement) mosquitoes should follow a hypergeometric distribution. Using the maximum likelihood estimate for the parameters of the hypergeometric distribution, a final estimate of ~12.6% in Murgon are direct migrants. The rate parameter of the Poisson processes controlling the intensity at which batch migration occurred was fit such that on average that number of mosquitoes migrated from Wondai to Murgon each month.

## Authors’ contribution

G.R. and B.J.T. designed research and obtained funding; J.M.M. obtained funding for simulation modeling; B.J.T. performed entomological surveys and sample collection; G.R. and I.F. generated and analyzed NGS data; S.L.W., T.M.L., J.B.B. performed simulation modeling with the input from G.R., H.C.S. and J.M.M.; I.F. and J.B.B. curated the data; G.R., I.F. and H.C.S. produced the figures; G.R. wrote the manuscript; all authors edited and approved the manuscript. The authors declare no competing interests.

## Acknowledgment

The research was funded by the research grant from a Mosquito and Arbovirus Research Committee (MARC) awarded to G.R. and B.J.T., a fund from the Commonwealth Scientific and Industrial Research Organisation (CSIRO) awarded to B.J.T to support entomological surveys and sample collection, research contract funds from the Australian Department of Health (DoH) awarded to the Mosquito Control Laboratory, and a DARPA Safe Genes Program Grant (HR0011-17-2-0047) awarded to J.M.M. We thank Greg Crisp and the team from the Wide Bay Public Health Unit, with assistance from Megan Nilon at the South Burnett Regional Council, Rachael Duncan and Glenn Proctor at the North Burnett Regional Council and Nadia Bannerman and Celia Dempster at the Gympie Regional Council. We thank Matthew Wessling and the team from the Central Queensland Public Health Unit for collecting mosquitoes in Central Queensland. We also thank Gregor J Devine and Brian J Johnson for insightful conversations on the control of *Ae. aegypti* in Australia.

## Supplemental Tables

**Table S1.** Results from the surveillance activities in the Wide Bay Burnett region (WBBR) over 4-10 weeks in 2018-19, and the survey years since 2004 with *Aedes aegypti* presence/absence.

| Town | % positive premises ($n_p$) | % positive ovitraps ($n_o$) | Weeks of trapping | Years present since 2004 | Years absent since 2004 |
| --- | --- | --- | --- | --- | --- |
| Wondai | 40 (10) | 20 (40) | 10 | 2013,-17,-18 | 2004,-14 |
| Murgon | 29 (7) | 57 (28) | 10 | 2013,-17,-18 | 2006 |
| Goomeri | 57 (7) | 32 (28) | 9 | 2004,-14,-18 | 0 |
| Gayndah | 60 (5) | 20 (20) | 5 | 2004,-14,-18 | 0 |
| Mundubbera | 100 (5) | 100 (24) | 4 | 2011,-12,-18 | 0 |
| Biggenden | 60 (5) | 100 (23) | 6 | 2011,-12,-18 | 0 |
| Gin Gin | 0 (8) | 0 (32) | 10 | 2011-17 | 2004,-18,-19 |
| Monto | 100 (5) | 100 (36) | 6 | 2011,-12,-18 | 0 |

**Table S2.** Average mosquito life cycle parameters. Time-varying parameters were scaled such that their mean values over the simulation period correspond to the values in this table.

| Parameter | Value | Reference |
| --- | --- | --- |
| Egg Duration (days) | 5 | Christophers, S. & Others. *Aedes aegypti* (L.) the yellow fever mosquito: its life history, bionomics and structure. *Aedes aegypti* (L.) the Yellow Fever Mosquito: its Life History, Bionomics and Structure. (1960). |
| Larval Duration (days) | 6 | Christophers, S. & Others. *Aedes aegypti* (L.) the yellow fever mosquito: its life history, bionomics and structure. *Aedes aegypti* (L.) the Yellow Fever Mosquito: its Life History, Bionomics and Structure. (1960). |
| Pupal Duration (days) | 4 | Christophers, S. & Others. *Aedes aegypti* (L.) the yellow fever mosquito: its life history, bionomics and structure. *Aedes aegypti* (L.) the Yellow Fever Mosquito: its Life History, Bionomics and Structure. (1960). |
| Egg production per female (day<sup>-1</sup>) | 20 | Otero, M., Solari, H. G. & Schweigmann, N. A Stochastic Population Dynamics Model for *Aedes Aegypti*: Formulation and Application to a City with Temperate Climate. *Bull. Math. Biol.* 68, 1945–1974 (2006). |
| Daily population growth rate (day<sup>-1</sup>) | 1.175 | Simoy, M. I., Simoy, M. V. & Canziani, G. A. The effect of temperature on the population dynamics of *Aedes aegypti*. *Ecol. Modell.* 314, 100–110 (2015). |
| Daily adult mortality rate (day<sup>-1</sup>) | 0.09 | Focks, D. A., Haile, D. G., Daniels, E. & Mount, G. A. Dynamic life table model for *Aedes aegypti* (Diptera: Culicidae): analysis of the literature and model development. *J. Med. Entomol.* 30, 1003–1017 (1993).<br>Horsfall, W. R. Mosquitoes: Their Bionomics and Relation to Disease. (1972).<br>Fay, R. W. The biology and bionomics of *Aedes aegypti* in the laboratory. *Mosq. News* 24, 300–308 (1964). |

